# Mitochondrial genome editing: a possible interplay of atherosclerosis-associated mutation m.15059G>A with defective mitophagy

**DOI:** 10.1101/2023.04.21.537899

**Authors:** Victoria A. Khotina, Vladislav A. Kalmykov, Alexander D. Zhuravlev, Vasily V. Sinyov, Igor A. Sobenin, Alexander N. Orekhov, Vasily N. Sukhorukov

## Abstract

**Background:** It was assumed that the cause of chronic inflammation in atherosclerosis is a disturbance of the innate immunity response, caused, among other factors, by mitochondrial dysfunctions. It was also suggested that mitochondrial dysfunction may be caused by heteroplasmic mutations in mitochondrial DNA. The aim of this study was to evaluate the effect of the mitochondrial nonsense mutation m.15059G>A on cellular functions in atherosclerosis: lipoidosis, pro-inflammatory response, and mitophagy.

**Methods:** The human monocytic cell line THP-1 and cytoplasmic hybrid cell line TC-HSMAM1 were used. An original approach based on the CRISPR/Cas9 system was developed and used for the elimination of mtDNA copies carrying the m.15059G>A mutation in the *MT-CYB* gene. Using this approach, the Cas9-TC-HSMAM1 cells with an eliminated m.15059G>A mutation was obtained. The gene expression levels of genes encoding enzymes related to cholesterol metabolism were analyzed by quantitative RT-PCR. The evaluation of pro-inflammatory cytokine secretion was assessed using ELISA. Mitophagy in cells was detected using confocal microscopy.

**Results:** In contrast to intact TC-HSMAM1 cybrids, in Cas9-TC-HSMAM1 cells, incubation with atherogenic LDL led to a decrease in the expression of the gene encoding fatty acid synthase (*FASN*). It was found that TC-HSMAM1 cybrids are characterized by defective mitophagy and are also unable to reduce the production of pro-inflammatory cytokines (to form immune tolerance) in response to repeated LPS stimulation. Elimination of mtDNA carrying the m.15059G>A mutation led to the restoration of immune tolerance and activation of mitophagy in the studied cells.

**Conclusions:** The m.15059G>A mutation was found to be associated with defective mitophagy, immune tolerance, and impaired metabolism of intracellular lipids due to upregulation of the *FASN*. Thus, this mutation may play an important role in atherogenesis due to its contribution to the chronification of inflammation, which aggravates the progression of atherosclerosis.

## 1. INTRODUCTION

Atherosclerosis is a chronic disease characterized by inflammation localized within the walls of human arteries. The initial trigger for the development of atherosclerosis is the accumulation of intracellular lipids by intimal cells, which occurs as a result of phagocytosis of modified LDL particles ^1^. Moreover, phagocytosis of modified LDL can trigger a cellular pro-inflammatory response, which is an innate immune response ^2^. When disturbances occur in the innate immune response, it can result in chronic inflammation, which may ultimately lead to the development of an atherosclerotic lesion. Mitochondrial dysfunction is thought to be one of the causative factors of impaired innate immunity during atherosclerosis development. The presence of heteroplasmic mutations in mitochondrial DNA (mtDNA) can result in the appearance of dysfunctional mitochondria in intimal cells ^3^. The primary mechanism for the removal of damaged mitochondria in cells is through mitophagy, a process in which mitochondria fuse with lysosomes and undergo subsequent degradation ^4^. It is known that some mutations in mitochondrial DNA can lead to disruption of the mitophagy process ^5,6^. As a result of defective mitophagy, damaged mitochondria remain inside cells and can provoke a chronic immune response ^7^, which contributes to the development of an atherosclerotic lesion accompanied by plaque formation ^8^.

Our previous studies have identified several heteroplasmic mutations in mtDNA that are associated with atherosclerosis. These mutations were found to be present in atherosclerotic lesions within the human aortic intima ^9–11^. Specifically, mutations in the *MT-RNR1* (m.1555A>G), *MT-TL1* (m.3256C>T), *MT-TL2* (m.12315G>A), and *MT-CYB* (m.15059G>A) genes were established to be associated with the formation of lipofibrous plaques in the intima of the human aorta ^9^. We have created cytoplasmic hybrid (cybrid) cell lines containing mitochondrial mutations associated with atherosclerosis in order to study the possible association between mutations in the mitochondrial genome and manifestations of atherosclerosis at the cellular level (pro-inflammatory response, accumulation of intracellular lipids) ^12^. Using these cybrid cells, we have shown the negative effect of mitochondrial mutations associated with atherosclerosis on the ability of cells to activate mitophagy and reduce the production of pro-inflammatory cytokines (to form immune tolerance) in response to repeated LPS stimulation ^8^.

In this study, we aimed to investigate the effect of a mitochondrial mutation on the processes of lipoidosis and the immune response. Specifically, we focused on the role of the nonsense mutation m.15059G>A in the mitochondrial cytochrome B gene (*MT-CYB*). This mutation leads to the formation of a truncated cytochrome B protein that can affect mitochondrial function. To eliminate mtDNA copies carrying the m.15059G>A mutation, we developed an original approach based on the CRISPR/Cas9 system ^13–15^. Using this approach, we successfully removed the mtDNA, carrying this target mutation, and created cells that does not carry the m.15059G>A mutation (Cas9-TC-HSMAM1), based on the TC-HSMAM1 cybrid cell line. This made it possible to evaluate the effect of the m.15059G>A mutation on cellular metabolism and functions. It has been found that this mtDNA mutation may be the cause of defective mitophagy.

## 2. MATERIALS AND METHODS

### 2.1 Vector construction

The strategy of mtDNA elimination from cybrid cells using a specific mitochondria-targeted CRISPR/Cas9 system was used in accordance with recent studies ^16–18^. The MitoCas9 vector was constructed, containing the full coding sequence of Sniper-Cas9 (*Streptococcus pyogenes*) with flanking mitochondrial targeting sequence (MTS) of human *COX8A* (Cytochrome C Oxidase Subunit 8A) and the T7 promoter sequence. The Streptavidin-SpyTag sequence was added after the MTS site through the S1m aptamer for simultaneous delivery of Cas9 nuclease and sgRNA into mitochondria ^19^. To increase the efficiency and specificity of mtDNA cutting with the MitoCas9 vector, biotinylated sgRNA was used. Site-specific mutagenesis of Sniper-Cas9 was performed to increase its stability and specificity, as described ^20–23^. The mtDNA-encoded gene cytochrome b (*MT-CYB*) with the m.15059G>A mutation was selected as a target for the MitoCas9 RNA complex.

### 2.2 Cell culture and transfection

The human monocytic cell line THP-1, as well as cybrid cell line TC-HSMAM1 and Cas9-TC-HSMAM1 cells were cultured in a complete culture medium of RPMI-1640 supplemented with L-glutamine, 10% (v/v) fetal bovine serum (FBS) (Gibco, USA), 100 U/ml of penicillin−streptomycin (Gibco, USA), and 50 mM β-mercaptoethanol (Sigma-Aldrich, USA) at 37 °C under a humidified atmosphere with 5% CO_2_. THP-1 was used as a control (reference) cell line. 2,5×10^5^ cells were seeded in a 24-well plate 24 h prior to the transfection. The MitoCas9 vector (1000 ng) was transfected by mannose-containing cationic liposomes in accordance with the N/P ratio of 6/1 (the ratio of the number of amino groups of the cationic lipid to the number of phosphate groups of the nucleic acid) ^24,25^. After 24 h the medium and the liposomes were removed by washing the cells twice with phosphate-buffered saline (PBS).

### 2.3 Analysis of the mitochondrial genome editing efficiency

For genome DNA extraction, Cas9-TC-HSMAM1 cells were lysed overnight at 60°C in SNET lysis buffer (20 mM Tris, pH 8.0; 5 mM EDTA, pH 8.0; 400 mM NaCl; 1% SDS) supplemented with Proteinase K (20 mg/mL) (Sigma Aldrich, USA). Genome DNA was purified using phenol:chloroform:isoamyl alcohol (25:24:1) according to standard protocol ^26^. The DNA concentrations were determined by the BioSpec-nano spectrophotometer (Shimadzu, Japan). Extracted genome DNA was used to amplify the surrounding region of the m.15059G>A mutation for T7 Endonuclease I (T7EI) mismatch detection assay using Q5 Hot Start High-Fidelity 2X Master Mix (New England BioLabs, USA) in accordance with recent studies ^27^. The PCR reaction was held in the thermal cycler T100 (BioRad, USA) in order to obtain PCR products. Purification and T7 Endonuclease I digestion of PCR products were made in accordance with the New England BioLabs protocol. Agarose gel electrophoresis of digested PCR products was held in the Sub Cell GT (BioRad, USA) horizontal electrophoresis system. Gel imaging was performed using the Gel Doc XR+ Gel Documentation System (BioRad, USA). To verify mtDNA cutting at the target site, we performed digital droplet polymerase chain reaction (ddPCR) on PCR products isolated from agarose gel. The amplification was conducted using a QX200 (BioRad, USA) and designed primers that detected double-strand DNA breaks (DSB) at the region of heteroduplex formation after Cas9 endonuclease cleavage of the gene sequence (**Table S1**). The number of events, which corresponded to the fluorescent response of a positive or cut mtDNA, and a negative template or wild-type mtDNA, was counted using QuantaSoft Analysis Pro software (BioRad, USA).

### 2.4 Preparation of human plasma low density lipoproteins

Plasma was derived from patients with atherosclerosis and myocardial ischemia, consistent with the World Medical Association (WMA) Declaration of Helsinki. Human low density lipoproteins (LDL) (d = 1.019-1.063 g/cm3) were isolated by preparative sequential ultracentrifugation using KBr for density adjustments, according to standard techniques ^28,29^. To prevent lipoprotein oxidation, all steps were performed in the presence of 3 mM EDTA. LDL preparations were dialyzed at 4°C in the dark against PBS containing 0.01 mM EDTA and sterilized through a polycarbonate filter with a 0.45 μm pore diameter (Corning, USA).

Protein content in LDL preparations was measured according to the method of Lowry et al. using Folin-Ciocalteu’s phenol reagent (Sigma-Aldrich, USA) ^30^. LDL preparations were stored in the dark at 4 °C for 7 days until use.

### 2.5 Generation of macrophage foam cells and measurement of their cellular cholesterol contents

THP-1, TC-HSMAM1 and Cas9-TC-HSMAM1 cells were maintained in a complete RPMI-1640 medium and 100 ng/mL of phorbol 12-myristate-13-acetate (PMA) (Sigma-Aldrich, USA) at 37°C for 48 h to induce cell differentiation to macrophages ^31^. Cell-derived macrophages were incubated in the presence of 100 μg/mL of previously isolated LDL for 24 h to induce lipid accumulation and foam cell formation. For the measurement of cellular cholesterol contents, the cells were lysed in a 0,2N sodium hydroxide solution, and the protein levels in the cellular lysates were then measured according to the method of Lowry et al., using 1 mg/mL BSA as a standard ^30^. Cellular lipids were extracted from cell samples using hexane–isopropanol (3:2; v/v). Cholesterol levels in the lipid extracts were measured using Fluitest CHOL (Analyticon, Germany) and a 1 mg/mL cholesterol solution as a standard. Total cholesterol content was calculated as the ratio of cholesterol to cell protein in each sample.

### 2.6 Analysis of lipid metabolism enzymes gene expression

Quantitative real-time PCR (RT-PCR) was used to analyze the gene expression levels of selected genes that encode enzymes involved in cholesterol metabolism in non-loaded and cholesterol-loaded macrophages.

Total cellular RNA was isolated using the ExtractRNA (Evrogen, Russia) with the removal of genomic DNA using the ezDNase Enzyme (Invitrogen, USA) according to the manufacturer’s protocol. RNA concentrations were determined by the BioSpec-nano spectrophotometer (Shimadzu, Japan). The RNA from each sample was then converted to cDNA using the MMLV RT kit (Evrogen, Russia) according to the manufacturer’s protocol. cDNAs from samples were amplified in triplicate with CFX96 Touch (BioRad, USA) using the qPCRmix-HS SYBR (Evrogen, Russia) for quantitative RT-PCR using designed primers (**Table S2**). The data were normalized relative to the median Ct values of endogenous controls (GAPDH and CAP1). The fold difference between the THP-1 control and treatments was calculated according to 2^-ΔΔCt^, where ΔΔCt=(ΔCt THP-1 control) - (ΔCt treatment) and ΔCt=(Ct target gene) - (the median Ct values of endogenous controls).

### 2.7 Evaluation of TNFα secretion of LPS-stimulated cells using ELISA

THP-1, TC-HSMAM1 and Cas9-TC-HSMAM1 cells were maintained in a complete RPMI-1640 medium supplemented with 10% FBS and 1 μg/mL lipopolysaccharide (LPS) from *Escherichia coli* O111:B4 (Sigma Aldrich, USA) at 37°C for 16 h (1^st^ LPS hit) to induce a pro-inflammatory response. After incubation, the samples were centrifuged at 200 g for 10 min and washed with PBS. The supernatant was removed, and fresh complete RPMI-1640 medium supplemented with 10% FBS with or without 1 μg/ml of LPS was added to the cells for additional incubation for 4 h (2^nd^ LPS hit). Then cell culture medium from each sample was aspirated and centrifuged at 200g for 10 min. The supernatants were used for subsequent analyses. The concentration of tumor necrosis factor α (TNFα) in supernatants was determined using TNFα ELISA Kit (R&D Systems, USA) and Nunc MaxiSorp ELISA Plates (Thermo Fisher Scientific, USA) following the manufacturer’s protocol. Absorbance was measured at 495 nm using the AMR-100T microplate reader (Allsheng, China).

### 2.8 Detection of mitophagy in cells using confocal microscopy

THP-1, TC-HSMAM1 and Cas9-TC-HSMAM1 cells were cultured in complete RPMI-1640 medium on a 6-well plate. Mitochondria were stained with 100 nM MitoTracker Green (Invitrogen, USA) for 30 min at room temperature. Then cells were incubated for 6 h at 37 °C under a humidified atmosphere with 5% CO_2_ in the presence of 5 μM carbonyl cyanide 4-(trifluoromethoxy)phenylhydrazone (FCCP) (Sigma-Aldrich, USA) to induce mitophagy. Then, lysosomes were stained with 50 nM LysoTracker Deep Red (Invitrogen, USA) for 30 min at room temperature. After that, cells were transferred to confocal dishes. The images were captured on a Leica DMi8 STELLARIS 5 confocal microscope (Leica Microsystems, Germany) with an HC PL APO CS2 63x/1.40 OIL objective.

### 2.9 Statistical analysis

All experiments were performed in triplicate to confirm the reproducibility of the results. IBM SPSS Statistics 27.0.1 software (IBM, USA) was used to analyze the data. All data were analyzed using the Student’s t test. Results of relative gene expressions and cytokine secretion are shown as the means ± SD. The captured confocal images were assessed using the CellProfiler software (Broad Institute of MIT and Harvard). Results of normalized mitophagy fold are shown as the means ± SEM. The level of significance was set to *p* < 0.05. Data visualizations for ddPCR analysis, relative gene expressions, cytokine secretion, and normalized mitophagy fold were prepared using GraphPad Prism 8 software (Graph Pad Software Inc, USA).

## 3. RESULTS

### 3.1 Elimination of m.15059G>A mutation in MT-CYB gene by MitoCas9 RNA complex

We have developed a method to eliminate the m.15059G>A mutation from the mitochondrial genome using the MitoCas9 RNA complex to target and remove all mtDNA copies carrying this mutation (**Figure 1, A**). The efficiency of cell transfection and elimination of mtDNA containing the m.15059G>A mutation was assessed using ddPCR with primers that detected DSB at the region of heteroduplex formation after cutting the gene sequence by Cas9 endonuclease. The cutting of mtDNA at the site of the m.15059G>A mutation was confirmed using the T7EI mismatch detection assay (**Figure 1, B**). As a result, the heteroplasmy level of the m.15059G>A mutation was reduced from 79.29 ± 11.58% to 12.81 ± 3.24% (p<0.001) in TC-HSMAM1 cybrids treated with the MitoCas9 RNA complex (**Figure 1, C**).

**Figure 1.**
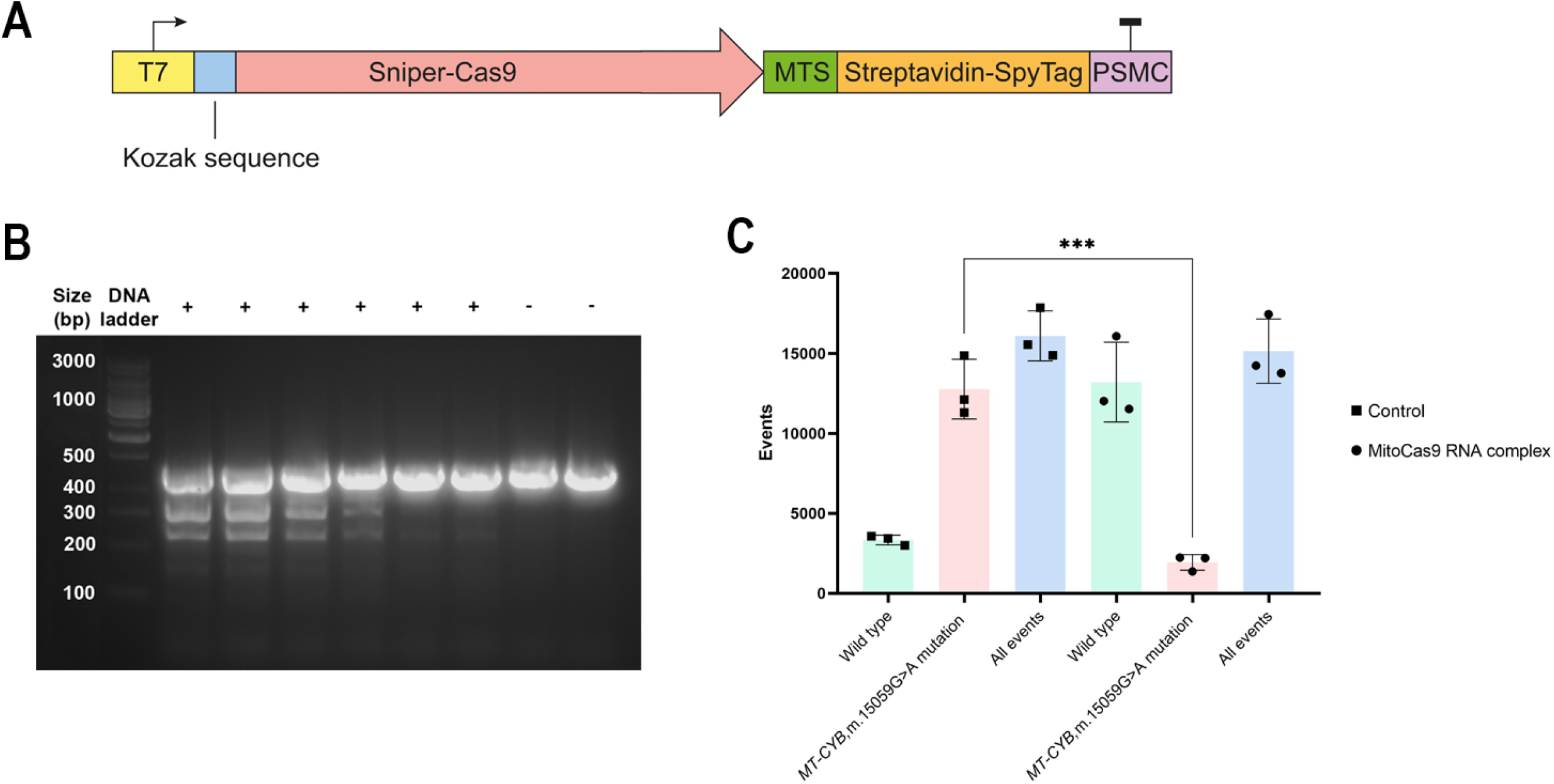
Elimination of m.15059G>A mutation in *MT-CYB* gene by MitoCas9 RNA complex: **A**. The MitoCas9 vector structure. T7 – T7 promoter sequence. Sniper-Cas9 – Cas9 coding sequence. MTS – mitochondrial targeting sequence derived from the *COX8A* (Cytochrome C Oxidase Subunit 8A) sequence. Streptavidin-SpyTag – streptavidin/biotin. PSMC – terminator region. **B**. Representative gel image of T7EI-treated PCR products amplified from the mtDNA with the m.15059G>A mutation. The T7EI method showed that heteroduplexes were identified in the six elevated regions by PCR reaction, indicating that the MitoCas9 RNA complex cleaved the double-stranded mtDNA at the target site. Moreover, it was showed that no T7EI cleavage was observed in two negative controls with PCR products amplified from cell mtDNA, where the MitoCas9 RNA complex was not applied. (+) PCR products amplified from the mtDNA with the m.15059G>A mutation, where the cleavage was found. (–) PCR products amplified from the wild type mtDNA, where the cleavage was not found. **C**. Efficacy of elimination of the mtDNA with the m.15059G>A mutation by the MitoCas9 RNA complex. Wild type – intact mtDNA without mutation. *MT-CYB*, mutation m.15059G>A – mtDNA carrying mutation. All events – the total number of mtDNA molecules. Control – TC-HSMAM1 cells without transfection with MitoCas9 RNA complex. MitoCas9 RNA complex – TC-HSMAM1 cells treated by the MitoCas9 RNA complex. Data information: *** *p*<0,001 (Student’s t test).

Thus, we have shown that the MitoCas9 RNA complex can effectively induce double-strand DNA breaks (DSBs) in the MT-CYB gene carrying the m.15059G>A pathogenic mutation in TC-HSMAM1 cytoplasmic hybrids. This approach allowed us to significantly reduce the heteroplasmy level of the m.15059G>A mutation to its minimum value.

### 3.2 Role of m.15059G>A in lipid metabolism

To assess the role of m.15059G>A in the intracellular cholesterol accumulation, THP-1, TC-HSMAM1 and Cas9-TC-HSMAM1 cells were incubated with LDL isolated from the blood of patients with atherosclerosis (atherogenic LDL). It was found that the initial content of total cholesterol did not differ significantly between all studied cell lines (**Figure 2A**). However, incubation of control THP-1 cells with LDL from atherosclerotic patients resulted in a 1.15-fold increase in cholesterol content (p<0.01), signifying the atherogenic properties of LDL in native cell culture. Furthermore, incubation of TC-HSMAM1 and Cas9-TC-HSMAM1 cells with atherogenic LDL resulted in a significant increase in total cholesterol levels by 1.29-fold (p<0.001) and 1.26-fold (p<0.001), respectively, when compared to cells not incubated with LDL. Nevertheless, there was no significant difference in total cholesterol values between the two cybrid cells that were incubated with atherogenic LDL. Since the elimination of mtDNA did not significantly affect the accumulation of cholesterol in cybrid cells, it suggests that the m.15059G>A mutation in the *MT-CYB* gene had no substantial effect on the intracellular cholesterol accumulation.

**Figure 2.**
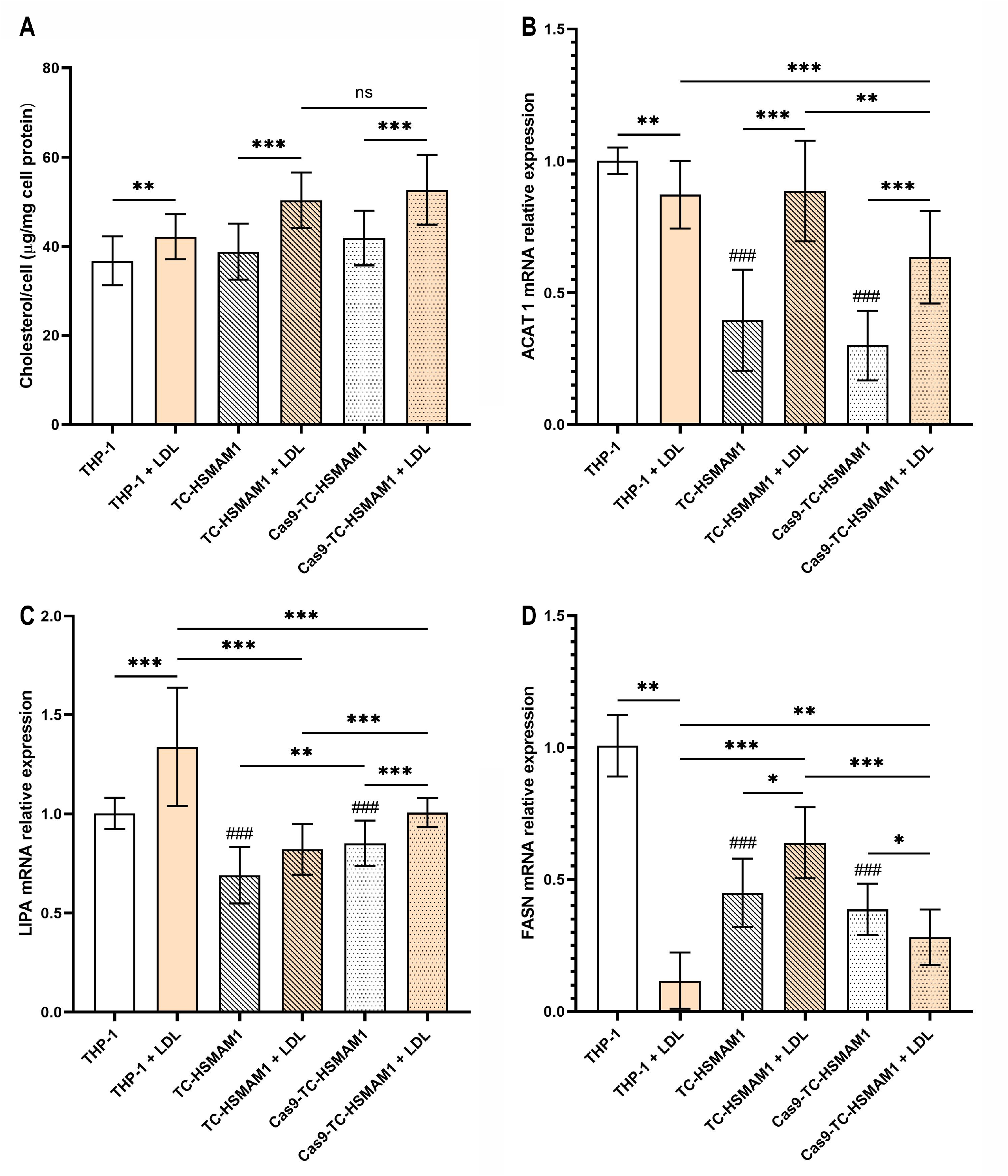
Evaluation of lipid metabolism in THP-1, TC-HSMAM1 and Cas9-TC-HSMAM1 cells: **A**. Total cholesterol levels of PMA differentiated THP-1, TC-HSMAM1 И Cas9-TC-HSMAM1 cells. **B**. *ACAT1* relative gene expression. **C**. *LIPA* relative gene expression. **D**. *FASN* relative gene expression. Data information: bars are mean ± SD. ns, not significant (*p*>0,05); * *p*<0,05; ** *p*<0,01; *** *p*<0,001; ### *p*<0,001 versus THP-1 (Student’s t test).

Intracellular lipid metabolism was assessed using a real-time PCR by the relative gene expression of the key enzymes of lipid metabolism: acetyl-CoA acetyltransferase 1 (*ACAT1*), which catalyzes the formation of acetoacetyl-CoA; lysosomal acid lipase (*LIPA*), also known as cholesterol ester hydrolase, which catalyzes the hydrolysis of cholesterol esters and triglycerides in lysosomes; fatty acid synthase (*FASN*), which catalyzes the synthesis of saturated fatty acids (**Figure 2, B, C, D**). It was found that the basal gene expression of *ACAT1, LIPA*, and *FASN* was downregulated in the TC-HSMAM1 and Cas9-TC-HSMAM1 cells compared to the control THP-1 cells (*p*<0.001). Additionally, there was no significant difference in the basal gene expression of the key lipid metabolism enzymes between the TC-HSMAM1 and Cas9-TC-HSMAM1 cells, with the exception of the *LIPA*, whose expression was increased in the Cas9-TC-HSMAM1 cells (*p*<0.01).

In THP-1 cells incubated with atherogenic LDL *LIPA* gene expression was increased (*p*<0.001), as well as *ACAT1* and *FASN* (*p*<0.01) gene expression was decreased, in comparison with the cells without incubation (basal gene expression). In TC-HSMAM1 cybrids treated with LDL, an increase of *ACAT1* (*p*<0.001) and *FASN* (*p*<0.05) gene expression compared to the basal expression was observed, while the expression of the *LIPA* gene remained unchanged. In Cas9-TC-HSMAM1 cells incubated with LDL, increased gene expression of *LIPA* and *ACAT1* (*p*<0.001) and decreased gene expression of *FASN* (*p*<0.05) were observed compared to the basal gene expression (**Figure 2, D**).

Taken together, the data on the expression of the key enzymes of lipid metabolism and the content of cholesterol in cybrids before and after the elimination of the proatherogenic mutation allow us to conclude that the m.15059G>A mutation can affect intracellular lipid metabolism by changing the expression of the *FASN* gene. It is interesting to note that the expression of this gene in Cas9-TC-HSMAM1 cells decreases in response to LDL, since the cells do not require *de novo* synthesized fatty acids in the presence of a source of exogenous fatty acids. However, a decrease in the expression of *FASN* was not observed in cybrids carrying the m.15059G>A mutation.

### 3.3 Innate immune tolerance of studied cells

The ability of intact THP-1, TC-HSMAM1 and Cas9-TC-HSMAM1 cells to form immune tolerance in response to pro-inflammatory stimulation with bacterial LPS was assessed by ELISA (**Figure 3**). Secretion of the pro-inflammatory cytokine TNFα was measured after the first (1^st^ LPS hit) and second stimulation (2^nd^ LPS hit) of cells with LPS.

**Figure 3.**
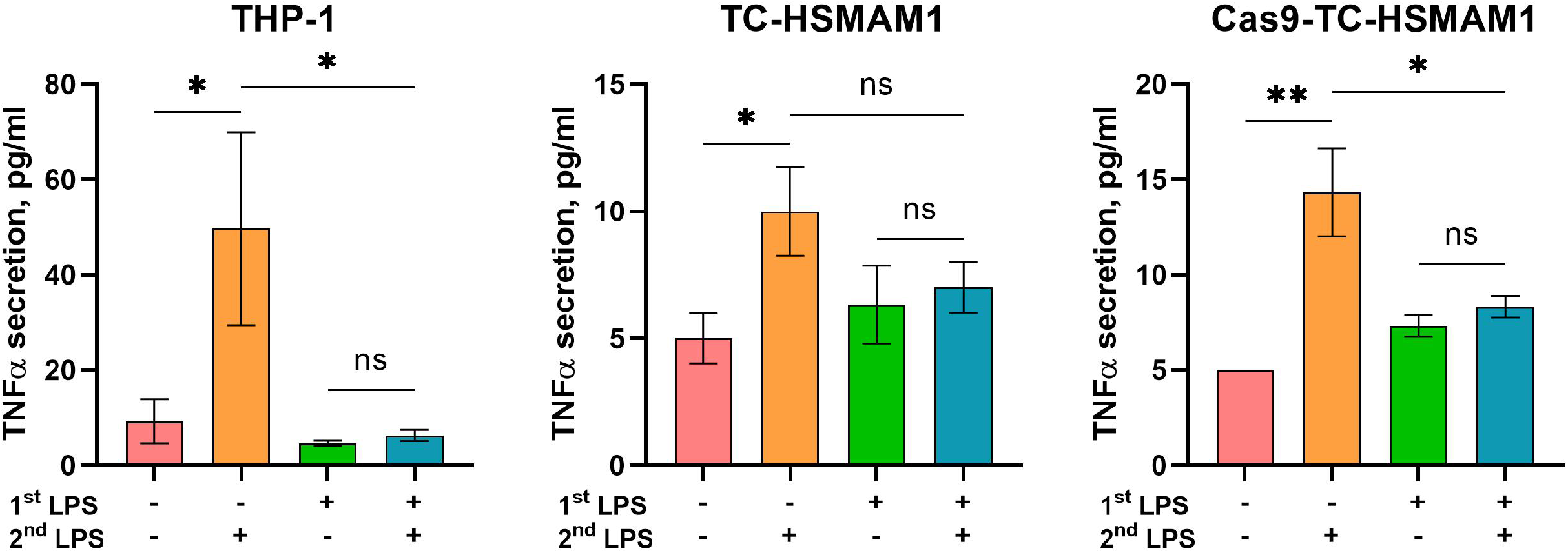
Innate immune tolerance of THP-1, TC-HSMAM1 and Cas9-TC-HSMAM1 cells: 1^st^ LPS – first LPS stimulation. 2^nd^ LPS – second LPS stimulation. Data information: bars are mean ± SD. ns, not significant (*p*>0,05); * *p*<0,05; ** *p*<0,01 (Student’s t test).

An increase in TNFα secretion in response to the first LPS stimulation and the absence of this effect after the second stimulation have been demonstrated in the control THP-1 cells (**Figure 3**). Thus, the control cells can form immune tolerance. At the same time, TC-HSMAM1 cybrids carrying the m.15059G>A mutation were found to have the inability to form immune tolerance in response to LPS stimulation. It was observed that these cells responded by TNFα secretion upon repeated LPS stimulation. However, TNFα secretion after second LPS stimulation was reduced (*p*<0.05) in Cas9-TC-HSMAM1 cells compared to the TNFα secretion after first LPS stimulation. Co, it was shown restoration of immune tolerance in Cas9-TC-HSMAM1 cells. Thus, the studied m.15059G>A mutation was found to have a negative effect on the immune tolerance of monocytes, preventing the resolution of inflammation.

### 3.4 Role of m.15059G>A mutation in mitophagy

Mitophagy activation in the control THP-1 cells, TC-HSMAM1 cybrids and Cas9-TC-HSMAM1 cells was assessed by the colocalization of mitochondria and lysosomes using confocal microscopy in the presence of MitoTracker Green and LysoTracker Deep Red. We assessed PINK-dependent mitophagy induced by depolarization of the mitochondrial membrane with FCCP in comparison to the basal level in unstimulated cells. In TC-HSMAM1 cells, FCCP did not induce a significant change in the level of mitophagy (**Figure 4**), indicating defective mitophagy in this cell line.

**Figure 4.**
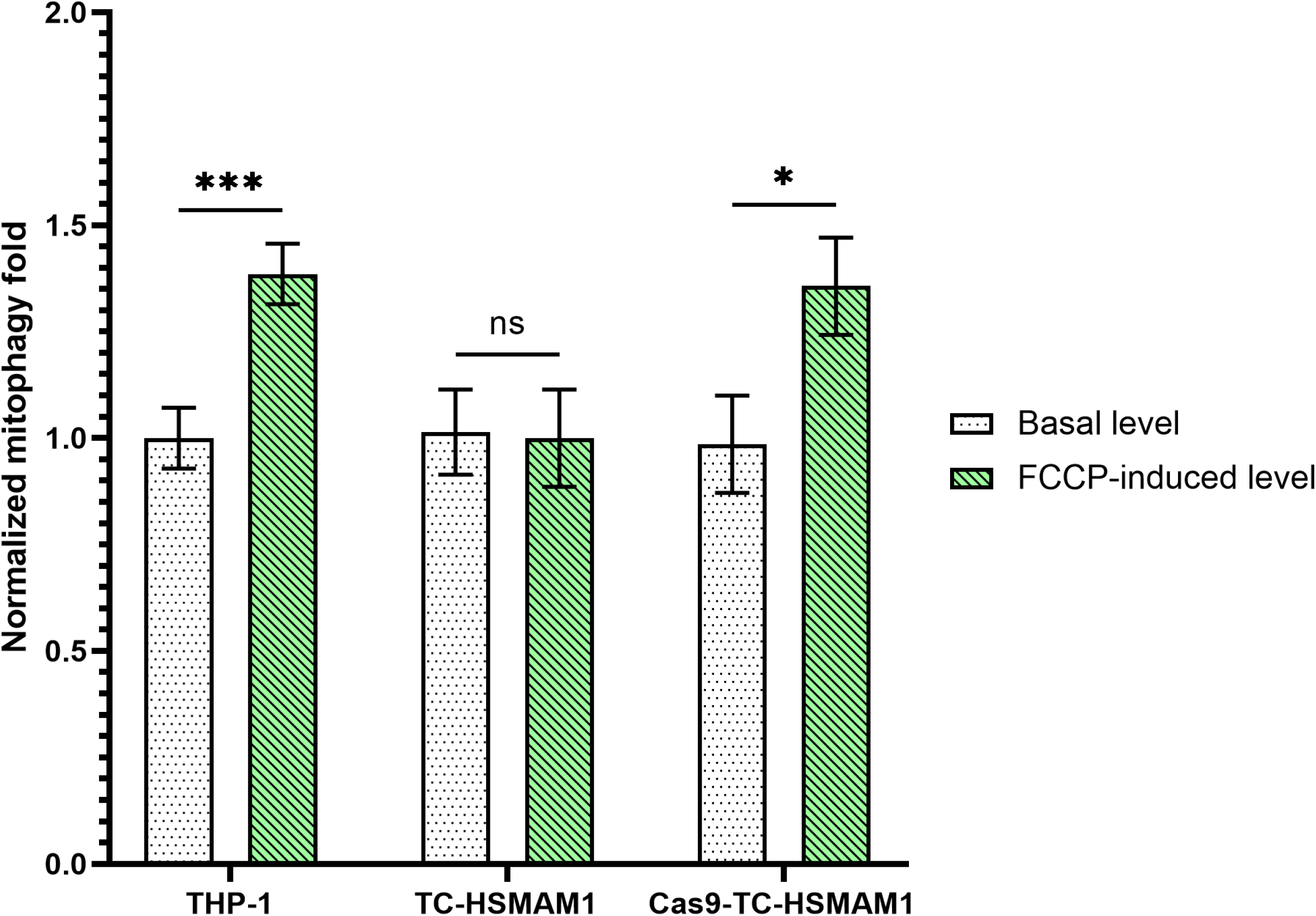
Normalized mitophagy fold in THP-1, TC-HSMAM1 and Cas9-TC-HSMAM1 cells: Level of mitophagy is the number of mitochondria colocalized with lysosomes relative to all mitochondria. Data information: bars are mean ± SEM. ns, not significant (*p*>0,05), * *p*<0,05, *** *p*<0,001 (Student’s t test).

In contrast, the level of mitophagy was increased by 1.39-fold (p<0.001) in FCCP-treated control THP-1 cells (**Figure 4**). Similarly, in Cas9-TC-HSMAM1 cells incubated with FCCP, the level of mitophagy was enhanced by 1.38-fold compared to the basal level (p<0.05). Confocal images accompanying the data on mitophagy activation demonstrated the colocalization of mitochondria with lysosomes, providing a comprehensive view of mitophagy dynamics (**Figure 5**).

**Figure 5.**
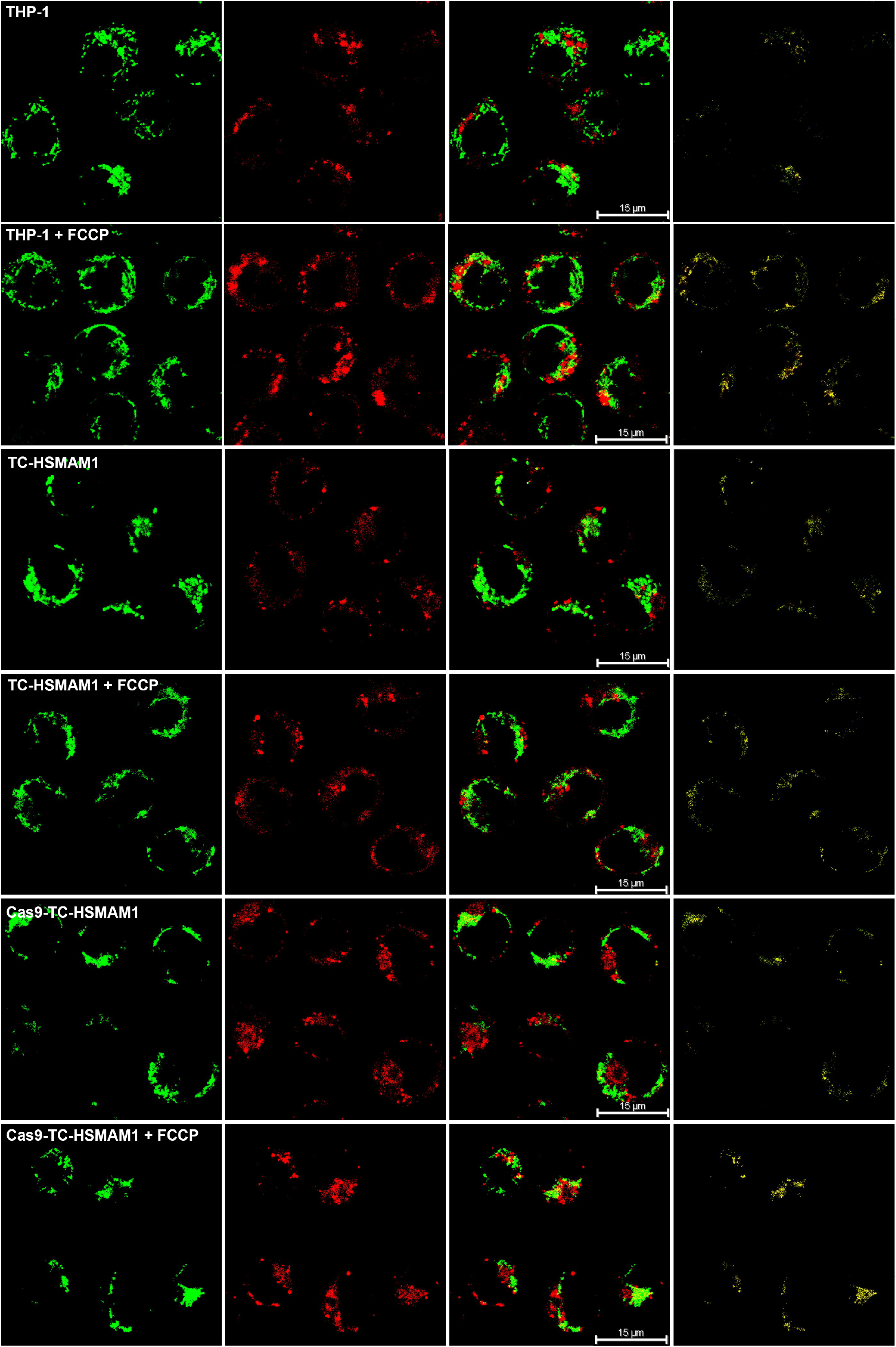
Confocal imaging of mitophagy in THP-1, TC-HSMAM1 and Cas9-TC-HSMAM1 cells. Mitochondria were stained with MitoTracker Green dye (green fluorescence), and lysosomes were stained with LysoTracker Deep Red dye (red fluorescence). Channel combination shows the colocalization of lysosomes with mitochondria. Lysosome-enwrapped mitochondria showed as yellow. Magnification 63x.

According to our results, the m.15059G>A mutation may be associated with the disruption of mitophagy processes in monocyte cells. Defective mitophagy, in turn, can lead to the preservation and accumulation of dysfunctional mitochondria in cells, and the aggravation of atherosclerosis.

## 4. DISCUSSION

The present study successfully demonstrated the effectiveness of the developed MitoCas9 vector in eliminating mtDNA copies containing the m.15059G>A mutation in the MT-CYB gene. Our findings are corroborated by previously reported data from studies utilizing CRISPR/Cas9-based mitochondrial genome editing systems to decrease targeted mtDNA copy number and levels of its transcripts ^16,18,32^. Our approach to eliminate mtDNA carrying the m.15059G>A mutation in the MT-CYB gene resulted in a significant reduction in heteroplasmy for this mutation. This allowed us to evaluate the effect of this mutation on the phenotypic and functional properties of the cybrid cells. In addition to the TC-HSMAM1 cybrids, we utilized the human monocytic cell line THP-1 as a reference for evaluating the impact of mitochondrial mutations on the functions of cybrid cells obtained from control THP-1 cells. While THP-1 cells and TC-HSMAM1 cybrids have the same nuclear genome, the original cell line cannot serve as a suitable control due to the presence of mitochondria with mtDNA from patients with atherosclerosis, which differs in genetic material from the mitochondria of the THP-1 cell line ^12^. Therefore, we evaluated the effect of the elimination of mtDNA with an atherogenic mutation on Cas9-TC-HSMAM1 cells and used the THP-1 cell line as a reference for the macrophage phenotype in a normal physiological condition.

In this study, we used the THP-1 cell line, as well as the TC-HSMAM1 cybrid cell line and Cas9-TC-HSMAM1 cells, to assess cell parameters such as the ability to accumulate intracellular lipids, produce pro-inflammatory cytokines in response to lipopolysaccharide stimulation, and the activation of mitophagy.

It was established that the examined mitochondrial mutation had no significant effect on the accumulation of intracellular cholesterol during the incubation of the studied cells with atherogenic LDL. The gene expression of *FASN* was found to be downregulated in Cas9-TC-HSMAM1 cells, indicating the impact of the m.15059G>A mutation on the expression of this gene in TC-HSMAM1 cybrids. Therefore, it can be concluded that the m.15059G>A mutation may be associated with a change in the intracellular synthesis of fatty acids, despite the fact that the intracellular cholesterol content was not modified. Numerous studies suggest a strong correlation between fatty acid synthesis and the development of atherosclerosis ^33,34^.

Accordingly, it was reported that the downregulation of the *Fasn* gene in mouse macrophages can contribute to the reduction of atherosclerotic lesions. Moreover, in our previous study, we have shown the association of the m.15059G>A mutation with atherosclerotic lesions in the human aorta ^9^. It may partly support the observed correlations in TC-HSMAM1, which include the upregulation of *FASN* gene expression compared to both the reference THP-1 cells and the Cas9-TC-HSMAM1 cells. It is well known that at the initial stage of atherosclerotic lesion formation, the main role is played by the differentiation of macrophages into foam cells, due to disturbance of intracellular lipid metabolism ^35^. In turn, the formation of foam cells in atherosclerotic lesions may be closely associated with mitochondrial dysfunction, contributing to the disruption of intracellular lipid metabolism ^36^. In this regard, the presence of macrophages with atherogenic mtDNA mutations in atherosclerotic lesions can aggravate the development of atherosclerosis.

TC-HSMAM1 cybrids were found to be unable to form immune tolerance in response to repeated stimulation of cells with LPS. Accordingly, the phenomenon of immune tolerance is manifested in a decreased immune response of monocytes upon secondary LPS stimulation, leading to diminished cytokine production ^37,38^. In the present study, Cas9-TC-HSMAM1 cells, as well as cells of the original THP-1 cell line, showed an adequate response to LPS stimulation, which consisted in an increase in TNFα secretion after the first LPS stimulation and a decrease in the secretion of pro-inflammatory cytokine after repeated LPS stimulation. The observed effect confirms the manifestation of immune tolerance in human monocytes. As a result, the m.15059G>A mutation may have a negative effect on the tolerance of monocytes, preventing the resolution of inflammation and contributing to its further chronification. Thus, the data obtained are consistent with our previous studies demonstrating the correlation of some heteroplasmic mtDNA mutations with pro-inflammatory activation of monocytes in patients with atherosclerosis ^39^.

Moreover, our results are consistent with the data on the possible association of pro-atherogenic mtDNA mutations with the inability of monocytes to develop immune tolerance ^8^.

We have found that the m.15059G>A mutation can negatively affect mitophagy activation in monocytes, leading to defective mitophagy. It is known that defective mitophagy, caused by the influence of certain agents on cells or mutations in nuclear genes, is accompanied by reduced activity of complex III of the mitochondrial respiratory chain ^40,41^. We hypothesize that in our study, an opposite effect was observed, where the impaired activity of complex III contributed to the reduction of mitophagy activity in cells.

Additionally, evidence of a link between mtDNA mutations and impaired processes of dysfunctional mitochondria disposal was demonstrated in a study where the m.12338T>C mutation in the mitochondrial gene *MT-ND5* led to reduced activity of complex I and disruption of mitophagy in cells ^5^. Our data on the association of the m.15059G>A mutation with impaired utilization of defective mitochondria in monocytes was consistent with the results of the evaluation of the ability of the studied cells to form immune tolerance. Thus, we have received evidence for our suggestion about the relationship between disturbances in the immune response of cells and defective mitophagy. It should be pointed out that fatty acid synthase may also be involved in LPS-stimulated and Toll-like receptor 4 (TLR4)-mediated macrophage activation, leading to increased secretion of pro-inflammatory cytokines ^42^. Consequently, the increased expression of *FASN* may be one of the causes of the increased secretion of TNFα in response to repeated stimulation of cells with LPS, indicating the inability of cells carrying the m.15059G>A mutation to form immune tolerance.

It is well known that mitophagy is one of the key processes involved in controlling the quality and quantity of mitochondria in cells ^43^. In turn, the close interaction between mitophagy and innate immunity is important for the protective immune response of cells to infectious agents. In the case of exposure to pathogens and the development of inflammation, the response of innate immunity can affect the dynamics and homeostasis of mitochondria, as well as promote mitophagy activation ^44^. It is most likely that disruption of the processes of removal of dysfunctional mitochondria can lead to the accumulation of mtDNA mutations, impairment of cellular functional properties, and subsequent cell death. On the other hand, disruption of autophagy processes in cells, in particular mitophagy, leads to the accumulation of defective mitochondria accompanied by an increase in ROS generation. ^45^. Furthermore, defective mitophagy promotes hyperactivation of inflammatory pathways, including abnormal inflammasome activation followed by the development of chronic systemic inflammation, neurodegenerative, myopathic, cardiovascular, and autoimmune diseases ^4,46^. We have recently shown that modulation of mitophagy in human monocytes can influence the inflammatory response of innate immunity, leading to a decrease in the secretion of pro-inflammatory cytokines, which further highlights the close relationship between the degradation processes of defective mitochondria and the development of inflammation ^8^.

In summary, the m.15059G>A mutation has previously been shown to be associated with atherosclerotic lesions in the human aorta. In this study, we have determined that the m.15059G>A mutation may be associated with defective mitophagy and the development of chronic inflammation due to the inability of monocytes carrying the mutation to form immune tolerance, as well as with disrupted intracellular lipid metabolism via upregulation of *FASN* gene. It can be assumed that the studied mutation may play an important role in atherogenesis by potentially contributing to the chronicization of inflammation, aggravating the progression of atherosclerosis. The m.15059G>A mutation in the *MT-CYB* gene can be used as a novel genetic marker of predisposition to atherosclerosis in the early stages of the disease. In addition, this mutation may be a potential target for the development of anti-atherosclerotic drugs and therapies.

## ACKNOWLEDGMENTS

V.A. Khotina and V.N. Sukhorukov designed this study. V.A. Khotina, V.A. Kalmykov, A.D. Zhuravlev and V.V. Sinyov performed experiments. V.A. Khotina and V.N. Sukhorukov performed data analysis and interpretation. I.A. Sobenin and A.N. Orekhov supervised the project. V.A. Khotina and V.N. Sukhorukov prepared the original manuscript draft. I.A. Sobenin and A.N. Orekhov reviewed and edited the manuscript. All authors reviewed and approved the manuscript.

## SUPPLEMENTAL MATERIAL

Tables S1–S2

Major Resources Table

